# Enhancing Brain Age Prediction and Neurodegeneration Detection with Contrastive Learning on Regional Biomechanical Properties

**DOI:** 10.1101/2025.03.25.645330

**Authors:** J. Träuble, L.V. Hiscox, C. Johnson, A. Aviles-Rivero, C.B. Schönlieb, G.S. Kaminski Schierle

## Abstract

The aging process affects brain structure and function, yet its biomechanical properties remain underexplored. Magnetic Resonance Elastography (MRE) provides a unique perspective by mapping brain tissue stiffness and damping ratio, observables that correlate with age and disease. Using a self-supervised contrastive regression framework, we demonstrate that MRE surpasses conventional structural magnetic resonance imaging (MRI) in sensitivity. Specifically, stiffness captures Alzheimer’s disease (AD), while damping ratio detects subtle changes associated with mild cognitive impairment (MCI). Our regional analysis identifies deep brain structures, particularly the caudate and thalamus, as key biomarkers of aging. The greater age sensitivity of MRE translates to superior differentiation of AD and MCI from healthy individuals, pinpointing regions where significant biomechanical alterations occur, notably the thalamus in AD and hippocampus in MCI. Furthermore, our results reveal biomechanical alterations in cognitively healthy individuals whose aging profiles closely resemble patients with MCI and AD. These findings highlight MRE’s potential as a biomarker for early neurodegenerative changes, aiding dementia risk detection and early intervention.

## Introduction

Aging is a complex biological process that impacts the human brain at multiple levels, including structural and functional properties^1–3^. Over the past decades, neuroimaging techniques have offered valuable insights into these age-related changes, revealing declines in grey matter volume, white matter integrity, and functional connectivity^4–7^.

More recently, Magnetic Resonance Elastography (MRE) has emerged as a promising method for characterising the biomechanical properties of brain tissue, capturing microstructural properties of neural tissue^8^, which are relevant for brain aging and neurodegeneration^9^. Unlike conventional Magnetic Resonance Imaging (MRI), which provides anatomical images and is typically used to measure brain volumes, MRE non-invasively characterises the brain’s viscoelasticity by yielding quantitative maps of brain tissue shear stiffness, reflecting tissue composition, and damping ratio, which relates to cellular organisation. MRE involves a conventional MRI scanner but with an actuation system and specialized imaging pulse sequences to generate and track shear waves as they propagate through tissue, from which tissue mechanical properties are reconstructed via an inverse problem^10^. Recent studies indicate that mechanical brain properties exceed traditional MRI measures in sensitivity to age-related changes^9,11,12^. For instance, whole-brain stiffness has shown a sensitivity to aging-related softening at a rate three times greater than volumetric atrophy rates observed with MRI^13^. Furthermore, this sensitivity extends to neurodegenerative diseases such as Alzheimer’s disease (AD), Parkinson’s disease (PD), and frontotemporal dementia (FTD), where abnormal mechanical alterations in specific brain regions have been reported^14–16^. There are some regional MRE studies that have shown that aging effects are not uniform across the brain, with the frontal, temporal, and occipital lobes displaying distinct mechanical signatures of aging, while deeper brain structures show differential responses depending on disease state^17,18^. However, current MRE studies predominantly focus on whole-brain or region-wide averages, underutilising the detailed information available in biomechanical maps, which can be captured via nonlinear relationships at the voxel level. Unlocking this level of detail could provide a more refined understanding of localized mechanical changes associated with aging and neurodegeneration, potentially enabling earlier detection when interventions may still be effective.

Brain age estimation—the prediction of chronological age from neuroimaging data—has gained prominence as a means to assess deviations from normative aging trajectories^19^. This technique has been predominantly applied to structural MRI modalities^20^ but has also been extended to other imaging techniques such as functional MRI (fMRI) and diffusion MRI^21^. In the field of brain age estimation, different modelling approaches have been used, ranging from traditional statistical kernel methods such as Gaussian processes^22^ to deep learning models like convolutional neural networks (CNNs)^23^, and more recently, advanced self-supervised learning approaches^24^. By leveraging a relatively large dataset of healthy individuals to establish a normative aging trajectory, these models effectively mitigate class imbalance issues often found in classification tasks, where disease cohorts are typically underrepresented. Recently, Claros-Olivares et al.^25^ presented a first approach in applying MRE-based features for brain age prediction, demonstrating the utility of mechanical biomarkers using CNNs. However, this study applied a more conventional supervised approach, was limited to healthy aging cohorts, and did not explore applications in neurodegenerative diseases or detailed regional analyses.

In this study, we integrate MRE-derived mechanical properties of the brain into a self-supervised approach utilising a contrastive regression framework for brain age estimation, a method particularly well-suited for low-data regimes and non-uniform data distributions^26^. Our approach significantly enhances the sensitivity of MRE-based models and thus enables the detailed study of pathological aging in neurodegenerative cohorts. We demonstrate that MRE significantly outperforms MRI in brain age prediction and neurodegeneration detection, with stiffness capturing later-stage neurodegenerative changes in AD and damping ratio detecting early alterations in mild cognitive impairment (MCI). Extending our analysis to deep brain regions, we identify the caudate and thalamus as key age-sensitive structures, reinforcing their relevance in brain aging. We also demonstrate that the thalamus is predominantly affected in AD, while the hippocampus is most impacted in MCI. Furthermore, our framework identifies cognitively healthy individuals whose biomechanical aging profiles resemble patients with MCI or AD, highlighting the potential for MRE in detecting neurodegenerative changes early. These findings not only establish MRE-derived mechanical properties as robust biomarkers of age-related brain changes but also position MRE as one of the first methods to reliably and non-invasively predict the onset of neurodegeneration.

## Results

### Leveraging Contrastive Learning for MRE-Based Brain Age Estimation

We investigate the mechanical properties—stiffness (μ) and damping ratio (ξ) —by employing a self-supervised learning framework for brain age prediction, leveraging adaptive neighbourhood selection within contrastive regression tasks (Fig.1).

**Figure 1:**
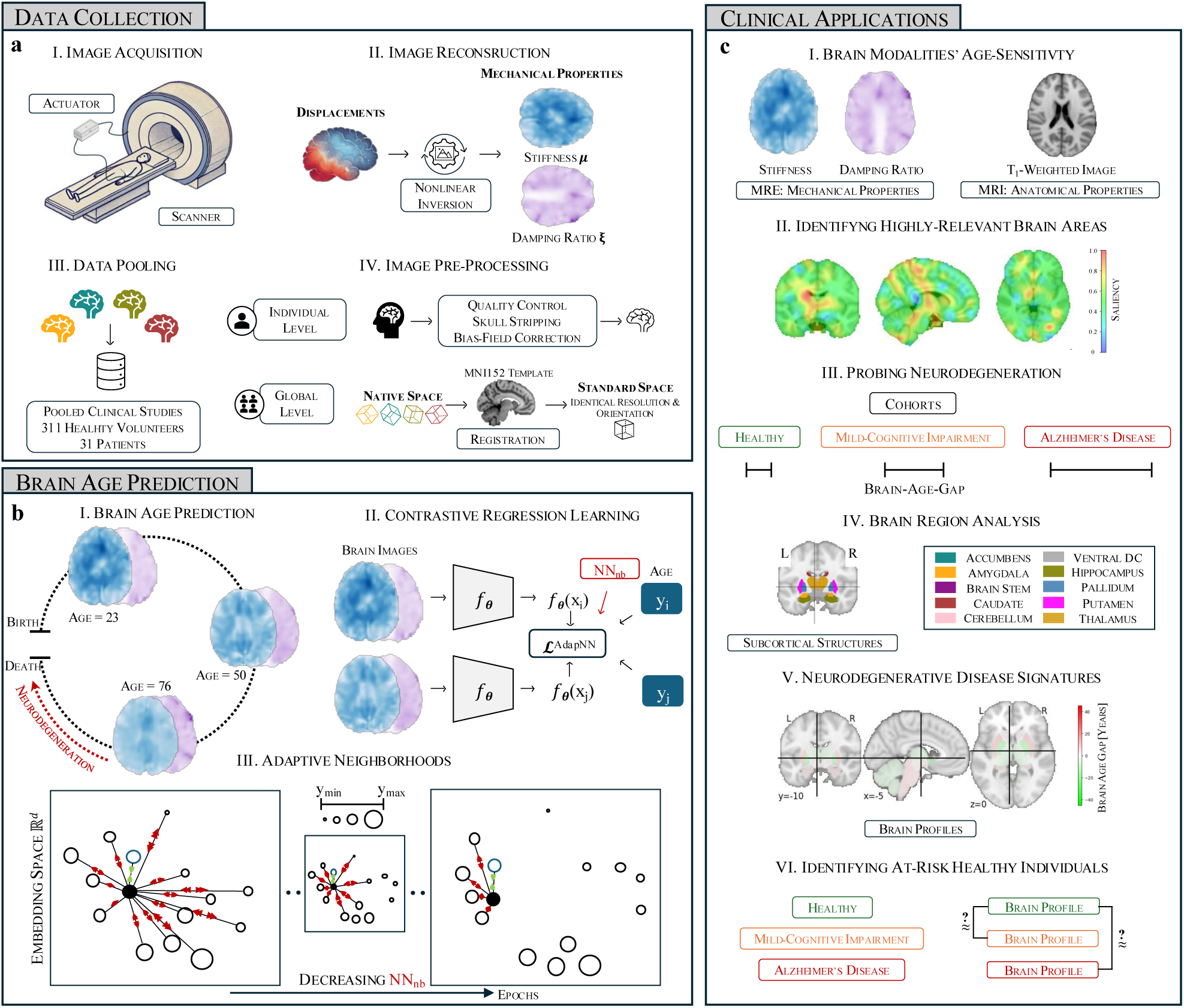
Study Workflow Overview. (a) Data collection and preprocessing: Image acquisition via MRE, reconstruction using nonlinear inversion to extract mechanical properties, data pooling from multiple clinical studies, and preprocessing steps such as skull stripping, bias-field correction, and registration to the MNI152 template. (b) Brain age prediction framework: A self-supervised contrastive regression model leveraging adaptive neighbourhood selection to enhance age-related feature learning. (c) Clinical applications: Predicting brain age trajectories during healthy aging to compare age-sensitivity of brain modalities, visualising most relevant brain areas, probing neurodegeneration using modelled normative aging trajectories, assessing regional differences across aging and disease, and identifying at risk healthy individuals showing similar brain profiles to disease signatures.

Our dataset comprises structural MRI and MRE data from 311 healthy volunteers and 31 patients, collected under closely matched acquisition protocols across multiple clinical studies. To the best of our knowledge, this represents the largest dataset of its kind. First, mechanical properties are extracted by applying nonlinear inversion techniques to brain tissue displacement data, resulting in quantitative maps of stiffness μ and damping ratio ξ^27^. Subsequent preprocessing—including skull stripping, bias-field correction, and registration to the MNI152 template—ensures that these maps are spatially standardized (Fig. 1a). These standardized maps serve as inputs for predictive modelling.

Central to our brain age prediction framework is the adaptive neighbourhood approach—a contrastive learning method tailored for regression tasks under non-uniform distributions in low-data regimes (Supplementary Fig. S2), which we have recently developed for stiffness maps^26^. In this work, we have extended this self-supervised framework to utilise additional neuroimaging data, such as damping ratio and MRI-derived anatomical images for comparison. Furthermore, we expand this framework to facilitate the integration of segmentation-based subcortical regions. Here, our contrastive learning method dynamically adjusts sample neighbourhoods to emphasize age-relevant differences (see Methods). The adaptive contrastive loss for a sample xi is defined as:

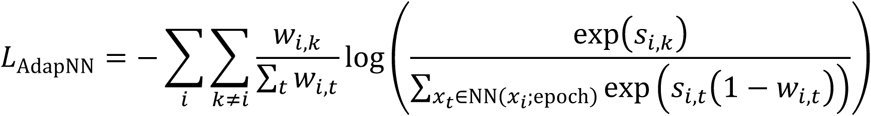

where wi,k = K(yi - yk) measures the similarity in age between samples xi and xk via a Gaussian kernel function K(·), and si,k = sim(f(xi), f(xk)) measures the similarity of their feature embeddings. The dynamically adjusted set NN(xi; epoch) ensures that the loss function focuses on the most relevant comparisons at each training stage. This allows the model to capture localised aging patterns and to generalize across the heterogenous dataset.

By integrating voxel-wise MRE-based measurements with advanced adaptive learning strategies, our approach (I) compares MRE to MRI in age sensitivity, (II) pinpoints brain areas highly relevant for predictions, (III) evaluates stiffness and damping ratio in neurodegeneration, (IV) analyses localised aging effects, (V) establishes neurodegenerative disease signatures in deep brain structures and (VI) identifies healthy individuals with signs of neurodegeneration.

### MRE outperforms MRI in Brain Age Estimation, with Brain Stiffness as the Dominant Aging Biomarker

We first examine the whole-brain average stiffness μ and damping ratio ξ across age. Global stiffness declines at a rate of -0.33% per year, highlighting progressive brain softening with aging (Fig. 2a) - consistent with trends observed in the field^28^. In contrast, the damping ratio increases at a rate of 0.34% per year, indicating more viscous or fluid-like tissue behaviour over time with higher energy dissipation (Fig. 2b). These trends indicate complementary aging trajectories. Representative mechanical brain maps at three different ages (Fig. 2c) illustrate these effects, showing a clear reduction in stiffness alongside the increase in damping properties. The strong association of both mechanical properties with age supports their suitability for brain age prediction. Building on these whole-brain trends, we utilise a voxel-wise approach to enhance resolution, allowing models to capture localized aging patterns across the brain.

**Figure 2:**
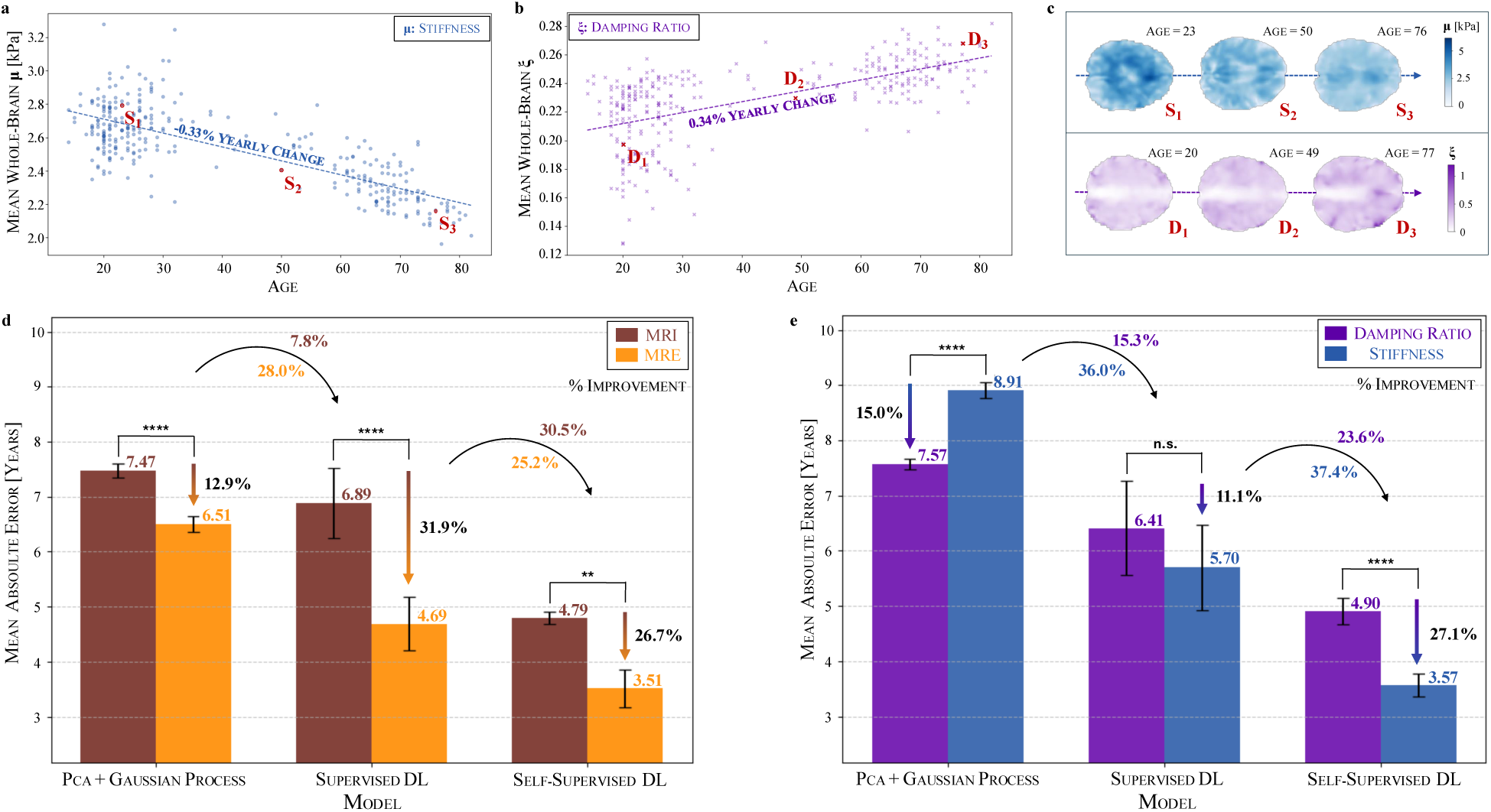
**Whole-brain biomechanical age trends reveal superior performance of voxel-wise MRE-based models over MRI in brain age prediction**. (a) Whole-brain average stiffness μ decreases with age, while (b) damping ratio ξ increases, reflecting distinct but complementary aging trajectories. (c) Representative mechanical property maps illustrate brain softening and increasing viscoelasticity across different ages. (d) Comparison of mechanical (MRE) against anatomical properties (MRI) using three distinct modelling approaches shows improved performance of MRE and that self-supervised deep learning achieves the lowest mean absolute error (MAE) for brain age prediction, outperforming PCA+Gaussian Processes and supervised deep learning. (e) Comparison of unimodal models (stiffness-only and damping ratio-only) highlights stiffness as the dominant mechanical biomarker of aging. N=10. n.s. (not significant) p ≥ 0.05; ** p < 0.01; **** p < 0.0001.

We compare three distinct modelling approaches for brain age prediction (Fig. 2d). The results demonstrate a significant improvement from established to more recent machine learning approaches, highlighting the impact of advanced modelling techniques on brain age prediction accuracy. While PCA with Gaussian processes achieves a median absolute error (MAE) of 7.47 years using MRI, deep learning improves prediction accuracy, with supervised deep learning reducing the MAE to 6.89 years and self-supervised learning further lowering it to 4.79 years. This trend is even more pronounced when incorporating MRE-based properties, where the MAE decreases from 6.51 years with PCA+GPs to 4.69 years with supervised deep learning, and further to 3.51 years with self-supervised learning. Notably, self-supervised learning further improves MRE-based predictions by 25.2% compared to supervised deep learning, highlighting the benefit of advanced representation learning for age prediction. Across all three model classes, the MAE is consistently lower when using mechanical brain properties derived from MRE compared to traditional T1-weighted MRI scans. For PCA+GPs, the difference between MRI and MRE is relatively small, with only a 12.9% reduction in MAE. However, for deep learning models, the advantage of MRE becomes much more pronounced, with supervised deep learning showing a 31.9% reduction in MAE and self-supervised learning achieving a similar 26.7% reduction. This superior performance of MRE-based models further underscores the higher age sensitivity of mechanical brain properties compared to structural MRI.

We now assess stiffness and damping ratio individually (Fig. 2e). While both mechanical properties capture aging-related changes, their effectiveness depends on the modelling approach. For kernel methods, damping ratio achieves a lower MAE (7.57 years) compared to stiffness (8.91 years), showing a 15.0% improvement. However, with more advanced models, stiffness emerges as the more informative feature for accurately modelling the aging trajectory. Supervised deep learning reduces the MAE to 6.41 years with damping ratio and further to 5.70 years with stiffness, a 11.1% improvement. With self-supervised learning, stiffness-based predictions achieve an MAE of 3.57 years, a substantial 27.1% improvement over damping ratio, reinforcing stiffness as the more informative feature. For all three methods, combining stiffness and damping ratio—as seen in the previous subfigure—yields better accuracy than either unimodal model. Remarkably, for self-supervised learning, stiffness alone (3.57 years) performs similarly to the combined MRE model (3.51 years), implying that, with sufficiently robust representation learning, stiffness alone captures most relevant age-related information.

We further investigate the spatial effects of aging (Supplementary Fig. S1) by normalising each image independently, removing the global trends of brain softening and increasing viscoelasticity observed in Fig.2 a–c. Under these spatially normalised conditions, MRE-based predictions remain highly effective, confirming that stiffness and damping ratio capture meaningful aging signals beyond global trends. This demonstrates that mechanical properties encode additional localised aging effects that persist independently of whole-brain trends, a novel finding that has not been previously reported and further highlights the unique sensitivity of MRE-derived biomarkers.

### Occlusion-Based Saliency Maps Reveal the Importance of Deep Brain Structures as Markers for Late Life Stages

To interpret and gain insights into the brain age prediction models, we apply occlusion-based saliency maps (see Methods) to examine which spatial features contribute most to the model predictions (Fig. 3).

**Figure 3:**
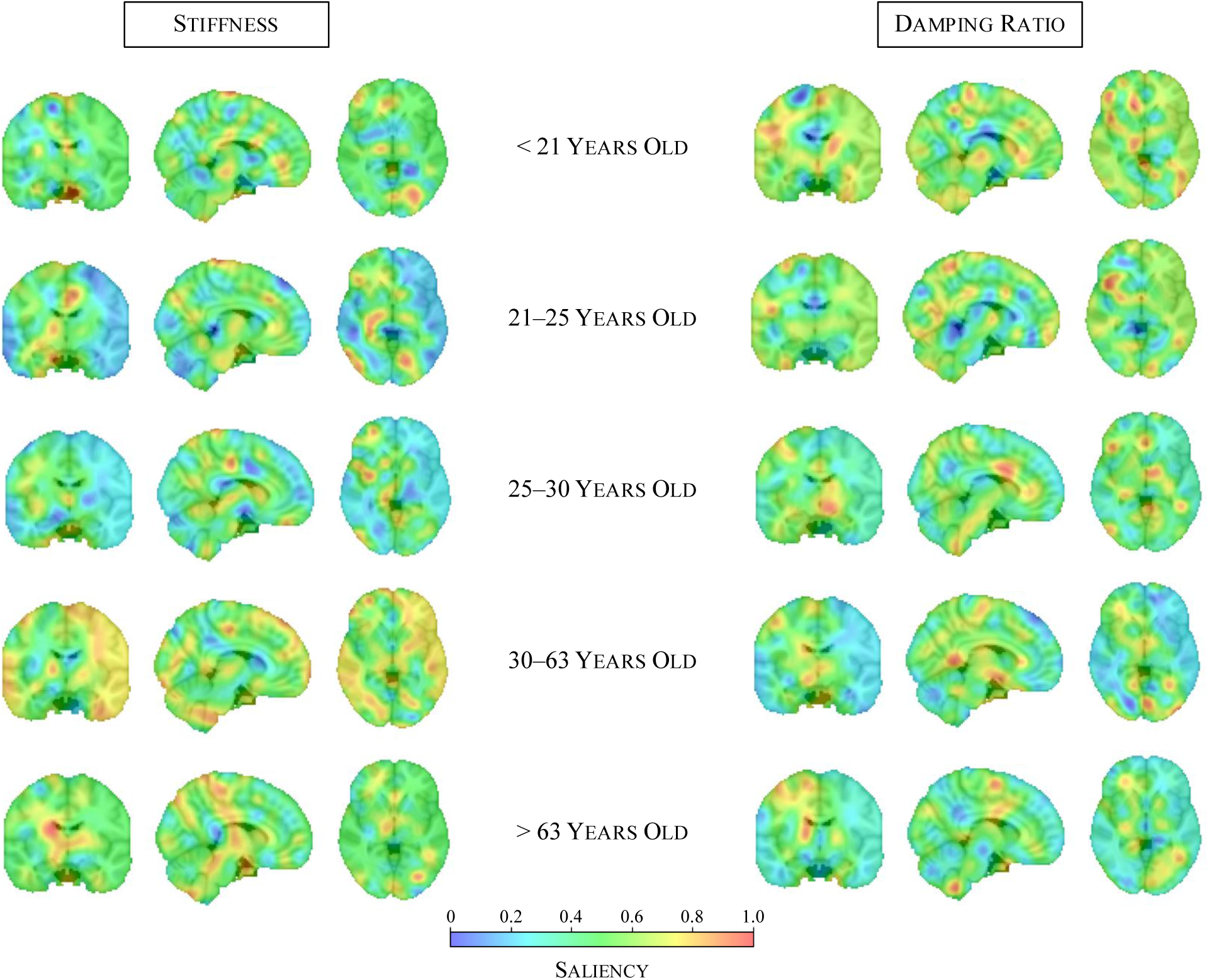
**Occlusion-based saliency maps highlight age-specific contributions of brain regions to brain age prediction**. Each row represents a different quantile-based age bin, illustrating the spatial distribution of important features for (left) stiffness-based models and (right) damping ratio-based models. Stiffness analysis reveals more focalised effects, with increasing cortical contributions in midlife and subcortical focus in older adults. Damping ratio exhibits a more diffuse distribution across the brain, suggesting broader viscoelastic aging effects.

The saliency maps reveal how the model focus shifts across the brain with age, reflecting changes in the spatial importance of mechanical properties for brain age prediction. At the same time, they highlight distinct patterns across biomechanical modalities. For the stiffness-based model, saliency appears more fragmented in younger age groups, with multiple small regions of high importance primarily in deeper brain structures, while some high-saliency areas also emerge near the cortical surface. In contrast, for midlife individuals (30–63 years), saliency becomes more spatially coherent, with a smoother distribution that is more prominently focused on cortical regions. This shows that stiffness alterations in cortical areas may be stronger indicators of aging during this period. In older age groups, the saliency distribution remains smooth rather than fragmented, in contrast to younger individuals. However, during older age, the model shifts its focus to subcortical structures, particularly the thalamus. Overall, deep grey matter structures, including the thalamus and putamen, consistently stand out across age groups, reinforcing their relevance in brain aging^18,29^. While the caudate, thalamus and putamen have previously been identified as the regions showing the most stiffness-related changes from childhood to adulthood^30^, our findings extend the importance of the thalamus in stiffness-related aging to later life stages.

For damping ratio-based predictions, the saliency maps reveal a more spatially diffuse pattern compared to stiffness, suggesting that damping ratio captures broader mechanical aging effects. In younger age groups, high-saliency regions are distributed across the brain, spanning both deep grey matter structures and more cortical regions. However, in midlife and older age groups, while the overall saliency distribution remains diffuse, contributions from cortical regions diminish, and the model increasingly focuses on deeper brain structures. Among these, the caudate continues to show strong contributions across all age groups. Our findings show that the importance of cortical grey matter does not extrapolate to later life, whereas the caudate remains a key region throughout aging. This extends previous studies^30^ that reported significant decreases in damping ratio within both the caudate and cortical grey matter during development from childhood to adulthood (ages 5 to 35 years), highlighting that while cortical effects are transient, the caudate’s relevance persists beyond early-life changes. The overall more diffuse saliency pattern indicates that viscoelastic aging effects may not be localized to specific structures but instead reflect more global mechanical alterations.

These findings reinforce that stiffness and damping ratio provide complementary information in brain age prediction, consistent with the improved performance observed in multi-modal models. While stiffness-based predictions exhibit a shifting spatial focus—initially more widespread in younger individuals, becoming more cortical in midlife, and then predominantly subcortical in older age—damping ratio-based predictions follow a more diffuse pattern, suggesting a more globally distributed role of viscoelastic changes in aging. The prominence of deep grey matter structures, particularly the thalamus, in both modalities highlight their fundamental role in brain aging, while the modality-specific spatial differences further support the idea that stiffness and damping ratio capture distinct but interrelated aspects of neurobiological aging.

### Superior Age Sensitivity of MRE Translates to Improved Detection of Disease Pathology of AD and MCI patients

We apply the self-supervised brain age models to cohorts of MCI and AD, and examine the brain age gap (BAG), i.e., the difference between predicted and chronological age, to assess whether the superior age sensitivity of MRE over MRI, observed earlier in healthy samples, translates into improved disease detection (Fig. 4a). Applying the MRI-based models to the MCI cohort, the predicted age distribution shows a similar but slightly decreased profile to the healthy samples (mean of BAG: -4.00 years, median of BAG: -6.83 years), showcasing slightly reduced brain ages in MCI samples. In contrast, MRE-based predictions show an increase in the brain age gap, with the median shifting from 0.04 years in healthy samples to 4.37 years in MCI samples, thus, observing an age acceleration effect. For the AD cohort, MRI-based predictions again show similar median values between healthy volunteers (7.85 years) and patients (8.10 years) with no detectable statistical significance between cohorts; however, the mean is elevated in the AD group (8.95 years) compared to healthy individuals (1.50 years), hinting at a slight age acceleration. In contrast, MRE-based predictions reveal a statistically significant increase in BAG, with the median rising from 1.96 years in healthy samples to 12.38 years in AD patients. These findings suggest that the greater age sensitivity of MRE over MRI observed in healthy aging encodes meaningful information for detecting early neurodegeneration, as it translates to superior differentiation in disease cohorts.

**Figure 4:**
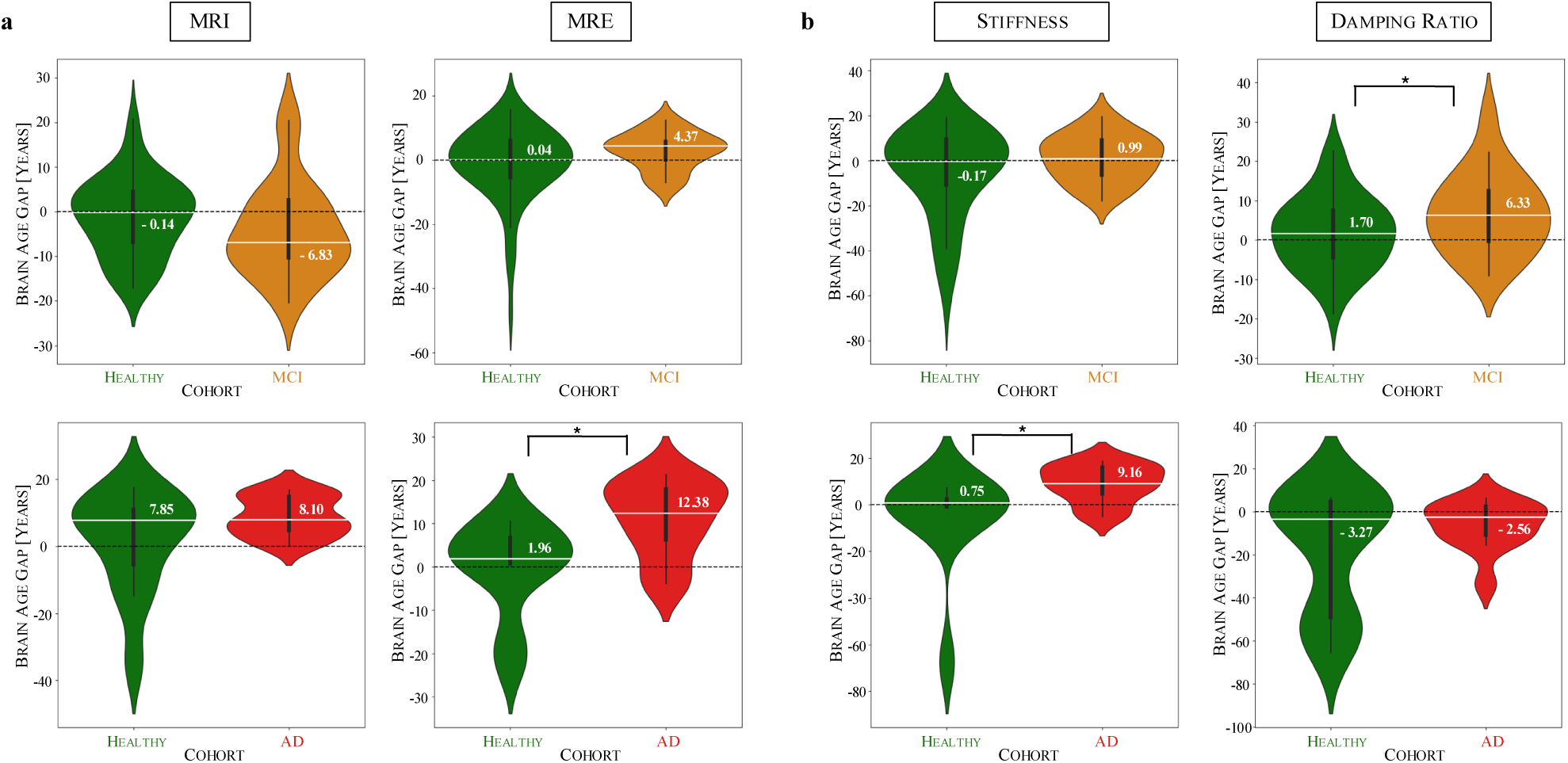
**Brain age gap (BAG) analysis highlights MRE’s sensitivity to detecting neurodegeneration-related defects in MCI and AD patients**. (a) MRI-based predictions show minimal differentiation in brain age gap (BAG) between healthy and diseased groups, while MRE-based models, using stiffness and damping ratio, capture age acceleration in MCI and AD. (b) Stiffness is more sensitive to AD-related changes, while damping ratio captures early-stage neurodegeneration-related defects in MCI. Median values are annotated. NMCI=10; NMCI control=68; NAD=11; NAD control=12. * p < 0.05.

To further dissect the individual contributions of stiffness and damping ratio to these effects, we examine their performance using stiffness-only and damping ratio-only models (Fig. 4b). In the MCI cohort, stiffness exhibits only a minimal increase in median BAG (from -0.17 to 0.99 years), while damping ratio shows a statistically significant increase (from 1.70 to 6.33 years), indicating that damping ratio predominantly drives the observed BAG elevation in MRE. Interestingly, the combined MRE model does not outperform damping ratio alone, implying that stiffness neither strongly reinforces nor counteracts the observed trend. In contrast, in AD, stiffness-based predictions show a significant increase in BAG (from 0.75 to 9.16 years), while damping ratio exhibits only a minor shift in median values. This suggests that stiffness is the dominant contributor to brain age acceleration in AD, while damping ratio adds auxiliary information that enhances the MRE-based model’s performance. Notably, unlike in MCI, the combined MRE model significantly outperforms damping ratio alone in AD, indicating that the integration of stiffness and damping ratio provides a more comprehensive characterisation of disease-related brain aging.

### Regional Brain Age Prediction Reveals Caudate and Thalamus as Key Age-Sensitive Structures

We extend the brain age modelling framework to individual brain regions, focusing on deep brain regions, which were identified as highly relevant in the whole-brain models through occlusion analysis earlier, as well as white matter (WM) and grey matter (GM), to investigate regional contributions to brain aging. Fig. 5a visualises the segmentation of the ten subcortical structures, illustrating the spatial distribution of the selected regions. The corresponding region sizes, shown in Fig. 5b, highlight the substantial variability in anatomical volume across structures, ranging from large regions such as the cerebellum (mean: 16,408 voxels) and thalamus (2,603 voxels) to smaller structures such as the nucleus accumbens (200 voxels) and the pallidum (633 voxels).

**Figure 5:**
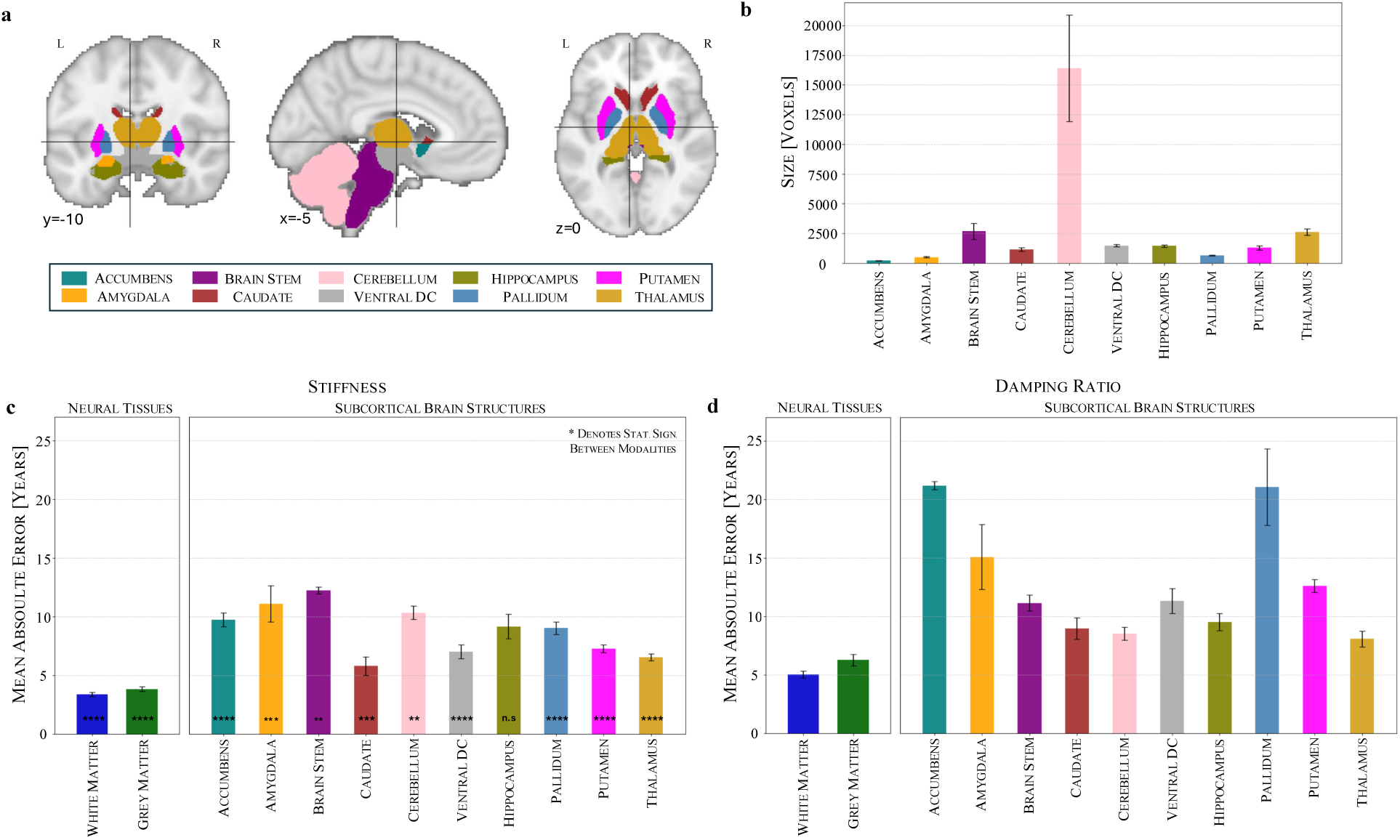
Subcortical structures, white matter and grey matter display distinct patterns in stiffness and damping ratio, and caudate and thalamus are key age-sensitive structures as revealed by brain age prediction errors. (a) Visualisation of ten subcortical structures used in the analysis. (b) Variability in anatomical volumes across regions. (c) Stiffness-based brain age prediction across subcortical regions shows caudate exhibits the lowest mean absolute error (MAE). (d) Damping ratio-based brain age prediction across subcortical regions shows thalamus displays the lowest MAE. N=10. n.s. (not significant) p ≥ 0.05; * p < 0.05; ** p < 0.01; *** p < 0.001; **** p < 0.0001.

Using the self-supervised learning framework, we assess region-wise brain age prediction separately for stiffness and damping ratio by training models using segmentation-based masks for each subcortical structure (see Fig. 5c and Fig. 5d). For both stiffness and damping ratio, GM and WM exhibit lower MAEs than any individual subcortical structure, likely due to their larger volume, which provides a more stable signal for model training. GM consistently shows higher MAEs than WM, with stiffness-based predictions yielding MAEs of 3.83 years for GM and 3.38 years for WM, while damping ratio-based predictions result in 6.27 years for GM and 5.03 years for WM. This finding may reflect differences in the underlying tissue composition and aging processes, as age-related changes in WM, such as demyelination and axonal degradation, are often more pronounced compared to GM. Furthermore, prior studies^31^ have shown that brain stiffness is correlated with myelin content, suggesting that changes in myelination could influence the predictive performance of stiffness-based models.

Among subcortical structures, the caudate achieves the lowest MAE for stiffness-based predictions (5.78 years), which is notable given its relatively small size. This is followed by the thalamus (6.54 years), ventral diencephalon (7.01 years) and putamen (7.26 years), while the pallidum (9.02 years) and hippocampus (9.15 years) perform moderately. The highest deviations are observed in the nucleus accumbens (9.75 years), amygdala (11.10 years) and brain stem (12.23 years), suggesting greater variability or reduced age sensitivity in these regions. For damping ratio-based predictions, the thalamus exhibits the lowest MAE (8.07 years), followed by the cerebellum (8.53 years) and caudate (8.97 years), suggesting that these structures may provide stable mechanical biomarkers of aging in the context of viscoelastic properties. The hippocampus (9.51 years), brain stem (11.15 years), ventral diencephalon (11.31 years) and putamen (12.60 years) show moderate errors, while the highest MAEs occur in the amygdala (15.07 years), pallidum (21.05 years), and nucleus accumbens (21.18 years). Comparing stiffness and damping ratio across structures reveals distinct trends, with caudate and thalamus consistently exhibiting lower errors for both stiffness and damping ratio, but other regions showing divergent behaviour, such as the pallidum and nucleus accumbens, which display much higher errors in damping ratio-based predictions compared to stiffness, suggesting reduced age sensitivity in viscoelastic properties. This suggests that stiffness and damping ratio encode complementary aspects of brain aging, supporting earlier findings that multi-modal models outperform unimodal models in capturing the full spectrum of age-related mechanical changes.

### Detecting Early Neurodegenerative Disease Signatures in Healthy Individuals and Identifying Thalamus and Hippocampus as Early Markers of AD and MCI, respectively

Having extended the brain age modelling framework to individual brain regions in healthy individuals, we now extend this regional analysis to disease cohorts to examine how neurodegenerative conditions affect mechanical brain aging (Fig. 6a). We compute brain age gaps (BAGs) for each region in MCI and AD cohorts, allowing us to construct subcortical brain age gap profiles for each group. To create a representative cohort profile, we average the individual brain profiles within each cohort.

**Figure 6:**
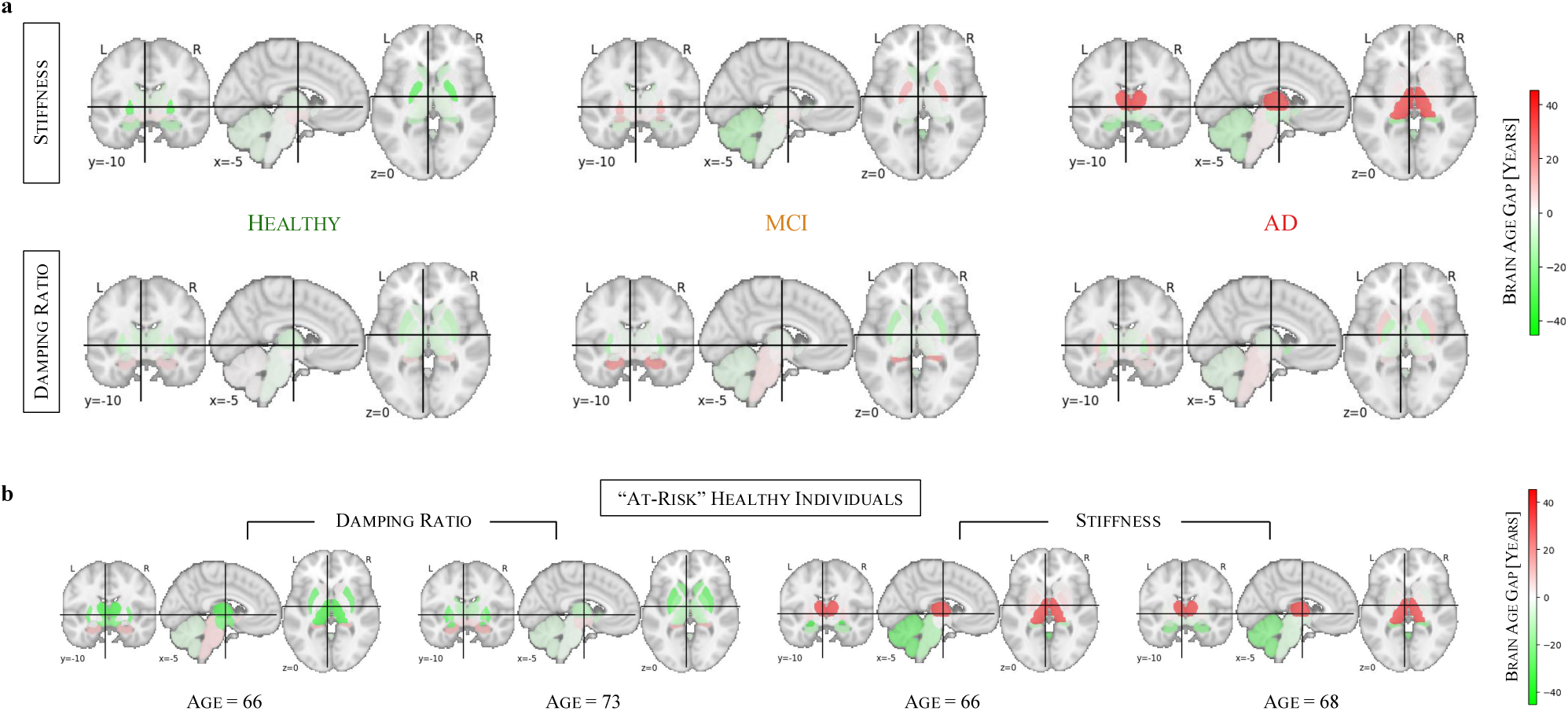
**Brain age profiles of disease cohorts and potentially at-risk subjects**. (a) Subcortical brain age gaps (BAGs) show distinct regional patterns in MCI and AD, with stiffness capturing AD-related changes and damping ratio highlighting early alterations in MCI. (b) Individual examples of healthy subjects whose brain age profiles resemble those of MCI or AD, suggesting potential early neurodegenerative changes.

In stiffness-based brain age profiles, healthy individuals show predominantly neutral to slightly negative BAGs across brain regions, suggesting that their predicted brain ages align closely with or are slightly younger than their chronological age. In the MCI cohort, some regions begin to exhibit moderately elevated BAGs, most notably in the pallidum, which shows the strongest increase, followed by the amygdala. This trend intensifies in the AD cohort, where the thalamus exhibits a markedly pronounced BAG elevation, while slight elevations are also observed in the pallidum and brain stem. The progression from healthy to MCI and AD is reflected in these brain age profiles, with increasing regional deviations marking the transition from normal aging to neurodegeneration. These findings mirror the whole-brain results, where stiffness has been the primary driver of brain age acceleration in AD. In damping ratio-based brain age profiles, healthy individuals similarly exhibit neutral to negative BAGs across most regions. However, in the MCI cohort, the hippocampus shows the strongest elevation, followed by the brain stem, while the ventral diencephalon exhibits only a slight increase, suggesting early-stage neurodegenerative changes. Unlike stiffness-based profiles, damping ratio-based predictions do not show a progressive increase in BAGs from MCI to AD. Instead, the AD cohort exhibits a brain profile similar to MCI, with the putamen showing the highest BAG elevation, while the hippocampus and brain stem remain elevated but without further progression, highlighting that viscoelastic changes predominantly reflect early neurodegenerative processes rather than later-stage disease progression.

Overall, these subcortical brain profiles provide regional insights into neurodegeneration and further support the differential contributions of stiffness and damping ratio to disease detection— stiffness being more sensitive to AD-related changes and damping ratio capturing early-stage alterations in MCI.

Beyond group-level analyses, these regional BAG profiles allow us to examine individual healthy participants and assess whether their brain age patterns resemble those observed in MCI or AD cohorts. Fig. 6b shows four such examples, where the first two healthy individuals are damping ratio-based profiles closely resembling the MCI cohort profile, while the latter two healthy individuals display stiffness-based profiles similar to the AD cohort. These similarities suggest that some healthy individuals may already exhibit early mechanical changes associated with neurodegeneration, potentially placing them at higher risk for future cognitive decline. This indicates that such individuals may be in a preclinical phase of AD, though further longitudinal studies are needed to confirm this.

## Discussion

Our comparative results show that MRE-based models consistently outperformed MRI-based models in brain age prediction, especially when using modern self-supervised learning techniques. This superior sensitivity to age-related change suggests that MRE can detect subtler brain aging effects than structural MRI. Unlike previous MRE studies constrained to regional averages, our voxel-wise deep learning approach captures complex, non-linear biomechanical dependencies, enabling us to move beyond global or regional stiffness and damping ratio trends. This granularity improves brain age prediction, with our contrastive regression framework outperforming kernel-based and supervised deep learning models. This highlights the importance of tailoring machine learning approaches to dataset characteristics, in our case, non-uniform, low-data regimes^32^. Our results reinforce the notion that stiffness is the dominant mechanical biomarker of brain aging and highlight the importance of using state-of-the-art modelling techniques to fully leverage MRE-derived properties. Our occlusion-based saliency analysis pinpoints regions with arbitrary shapes and sizes of most interest to the brain age models. Notably, the saliency maps reveal a shift in the most predictive brain regions across different age groups, supporting previous findings that mechanical aging patterns evolve throughout the lifespan^12,33,34^. Stiffness-based brain age models shift focus from cortical to subcortical regions, increasingly emphasising the thalamus and caudate in older adults, extending their known role in mechanical aging beyond early life^30^. Meanwhile, damping ratio-based models show a progressive decline in cortical relevance in older ages, revealing that cortical sensitivity to aging, previously observed only during the developmental years (ages 5-35)^30^, does not persist into later adulthood. Beyond healthy aging, MRE effectively detects neurodegenerative changes. While MRI-derived models show minimal BAG differences, MRE reveals significant deviations. Our results reveal that damping ratio is more sensitive to early MCI-related neurodegeneration, while stiffness more effectively captures age acceleration in AD.

This aligns with prior findings that damping ratio is strongly associated with cognitive function and poorer episodic memory performance, making it a marker of early cognitive decline^35,36^. In contrast, stiffness decline reflects advanced neurodegeneration, with significant reductions observed in AD patients^33,37^. These findings further reinforce the distinct pathological processes tracked by mechanical properties—damping ratio indicating early neurodegenerative disruptions and stiffness decline marking later-stage degeneration. Regionally, the caudate and thalamus emerge as the most age-sensitive subcortical structures. Previous MRE studies have primarily characterised the caudate and thalamus in terms of their age-related decline in stiffness, with an accelerated decrease observed in elderly individuals^38^. Our voxel-wise approach extends these findings by not only confirming their role in age-related stiffness changes but also demonstrating their relevance for damping ratio-based biomarkers. We further observe distinct regional prediction errors between both mechanical properties further suggesting that they capture different aspects of aging. Applying regional models to disease cohorts reveals that stiffness-based BAGs primarily detect age acceleration in AD, while damping ratio-based predictions highlight early neurodegenerative changes in MCI. Notably, some cognitively healthy individuals exhibit regional profiles resembling MCI or AD, suggesting MRE-based biomarkers may serve as early indicators of neurodegeneration.

In addition to previous findings on caudate atrophy and metabolic decline in aging and neurodegeneration^39,40^, our results emphasize the importance of biomechanical characterisation in detecting these processes. Studies have shown that caudate atrophy is associated with gait dysfunction and poorer physical performance^40^, while metabolic reductions in the caudate nucleus serve as sensitive biomarkers for normal aging and early neurodegenerative stages^39^. Furthermore, recent research highlights dopaminergic deficits in the caudate as a contributing factor to cognitive decline in Parkinson’s disease, particularly in pre-dementia stages^41^. Our findings extend this understanding by demonstrating that biomechanical measures offer a novel and sensitive marker of age-related changes in the caudate, reinforcing its role as a key region in neurodegeneration detection. Similarly, beyond well-documented atrophy and morphological alterations of the thalamus in neurodegenerative diseases^42,43^, our study highlights the potential of biomechanical properties as complementary biomarkers of aging and disease progression. Thalamic morphology has been proposed as a putative biomarker across multiple neurodegenerative disorders^42^, with distinct patterns of atrophy observed in early- and late-onset Alzheimer’s disease^43^. Our results build on this by showing that stiffness and damping ratio capture differential aging effects within thalamic subregions, further underscoring the relevance of MRE-derived biomarkers in tracking neurodegenerative progression.

Despite these advantages, limitations remain that provide avenues for future improvement. While our dataset is large for MRE studies, it remains smaller than MRI-based datasets^44^, which may explain the lack of statistical significance in MRI-based models for AD detection. However, the ability to detect significant effects in a relatively small MRE sample further supports its increased sensitivity, suggesting that MRE captures relevant biomechanical changes even with lower sample sizes. Furthermore, the inclusion of multiple clinical studies introduces potential confounding factors, which we mitigate through image registration, but some inter-study variability may persist. A further limitation is the need to retrain models without healthy controls when computing brain age gaps (BAG) for disease cohorts, making them susceptible to dataset biases. Larger and more diverse datasets will be crucial for improving generalizability. Additionally, current MRE studies occasionally exhibit partial brain coverage, which could affect regional analyses. Similarly, brain stem measurements remain challenging due to the attenuation of shear waves in centre of the brain along with its proximity to pulsating structures that can cause motion artifacts. Continued advancements in MRE hardware and sequence design will help mitigate these issues.

Our findings suggest promising directions for further research. Some “healthy” individuals exhibited biomechanical brain aging patterns similar to those seen in patients with MCI or AD. While these participants all scored within the normal range on the Montreal Cognitive Assessment (MoCA), standardised cognitive tests are not currently sensitive enough to identify participants that may be within the preclinical phase of disease. While this was beyond the scope of the current study, future large-scale longitudinal studies should consider whether our methods are able to prospectively identify participants that may go on to develop cognitive impairment and dementia. We now plan to apply the methods developed here to examine mechanical brain aging patterns in those at risk for AD, identified through genetics and other lifestyle factors, and integrate MRE with multimodal approaches—including other advanced MRI-based microstructural imaging techniques^45,46^—to improve sensitivity and enhance clinical applicability.

In summary, our findings establish MRE-derived mechanical properties as robust biomarkers of age-related brain changes and highlight MRE’s potential for non-invasive detection of biomechanical alterations associated with neurodegeneration.

## Data/code availability

The code is written in Python and PyTorch^47^ was used for model implementation. The code will be made publicly available on GitHub upon publication. Data from all studies can be accessed through a formal data-sharing agreement with CJ. Specifically, data from Study 1 and Study 2 are available at <u>MRE134</u> and <u>NITRC</u>, respectively.

## Methods

### Data

We compiled a dataset of 311 healthy individuals and 31 patients from multiple clinical studies. Each sample contains stiffness and damping ratio maps, as well as a T1-weighted anatomical scan. The healthy cohort was aggregated from five independent studies, each using highly similar MRE acquisition protocols and covering a broad age range (mean age: 41.0 ± 21.9 years). In addition, we assembled two disease cohorts comprising 31 patients, including 20 individuals classified with Mild-Cognitive Impairment^48^, and 11 individuals diagnosed with Alzheimer’s Disease^16^. All studies used a 3T MRI scanner, a Resoundant pneumatic actuator pillow system with a frequency of 50 Hz and a 3D spiral MRE sequence^49,50^. This dataset represents one of the largest collections of MRE-derived brain mechanical properties to date. Detailed information of our dataset can be found in Table 1.

**Table 1.** Detailed dataset information. *: This study is a pooled study itself.

|  | Study | Published In | #Subjects | Age [Years] | Sex [F:M] | Resolution [mm<sup>3</sup>] |
| --- | --- | --- | --- | --- | --- | --- |
| Healthy | 1* | 11 | 134 | 23.4 ± 4.0 | 78:56 | 2.0 |
|  | 2 | 51 | 60 | 37.8 ± 20.9 | 34:26 | 1.5 |
|  | 3 | 16 | 12 | 69.4 ± 2.4 | 6:6 | 1.6 |
|  | 4 | 48 | 68 | 69.3 ± 5.8 | 49:19 | 1.25 |
|  | 5 | 52,53 | 37 | 49.1 ± 16.6 | 16:21 | 1.25 |
|  | **Total** | - | **311** | **41.0 ± 21.9** | **183:128** | **2.0** |
| Disease | 6 | 16 | 11 | 76.8 ± 5.3 | 7:4 | 1.6 |
|  | 7 | 48 | 20 | 72.6 ± 8.5 | 14:6 | 1.25 |

### Image Pre-Processing

Displacement fields obtained from MRE acquisitions were processed using the nonlinear inversion (NLI) algorithm^27^, which estimates the complex shear modulus *G* ∗ = *G*′ + *iG*′′. Here, G’ represents the storage modulus, while G’’ corresponds to the loss modulus. From these values, we derived key mechanical properties: the stiffness measure μ as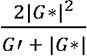 and the damping ratio ξ as 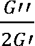^54,55^. These computed maps of stiffness and damping ratio were used as inputs for further analysis. Next, the MRE magnitude map and T1-weighted scans underwent skull stripping using FreeSurfer^56^ to extract brain tissue while removing non-brain structures. A bias field correction was then applied to eliminate intensity gradients that could introduce inconsistencies in the analyses. To mitigate inter-study variability, we performed affine registration of the images to the MNI152 template. This registration was carried out using ANTs^57^ at an isotropic resolution of 2 mm³, with final image dimensions of 91 × 109 × 91, ensuring uniform orientation and scale across all data. Finally, we normalised the quantitative stiffness and damping ratio maps by adjusting their mean to zero and standard deviation to one across the entire dataset. This normalisation approach preserves global trends of brain softening and increasing viscoelasticity with age and was consistently applied throughout the analysis. An exception was made for the analysis in Supplementary Fig. S1, where each image was independently normalised to a zero mean and unit standard deviation to specifically examine the spatial distribution of mechanical properties across aging. T1-weighted images were normalised on a per-image basis using the same standardization method. Furthermore, to account for variability in brain coverage within the quantitative mechanical property maps, T1-weighted images were masked accordingly to ensure consistent coverage across modalities.

### Contrastive Regression Framework for Brain Age Estimation

Typically, self-supervised and contrastive learning approaches are primarily designed for classification tasks, where samples are grouped into discrete categories. However, brain age prediction is a regression problem, requiring a fundamentally different formulation where similarity is determined by continuous age differences rather than class labels. Thus, we employ a contrastive regression framework designed to predict brain age from MRE-derived stiffness and damping ratio maps, as well as MRI images. This approach extends the previously developed Localized Neighbourhoods method^26^, which employs contrastive learning to adaptively adjust sample neighbourhoods in non-uniform regression tasks.

Unlike classification-based contrastive learning, where positive and negative pairs are predefined, our contrastive regression loss ensures that samples are attracted or repelled based on their age similarity:

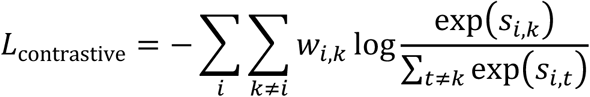

where:

- si,k is the cosine similarity between embeddings *f*(xi) and *f*(xk).
- wi,k is an age-aware weighting function, defining sample similarity based on age difference.

Following Barbano et al.^24^, we integrate:

Y-Aware weighting, where wi,k =K(yi−yk) controls sample attraction/repulsion strength using a Gaussian kernel:

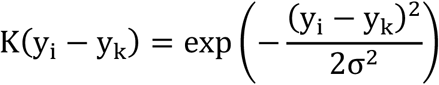

This ensures that age-similar samples are drawn closer while distant ones are repelled.

1. Exponential Scaling, modifying the denominator to increase repulsion strength for dissimilar samples:

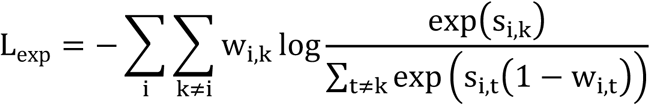

which adjust repulsion strength based on the samples’ age difference.

Further, following the Adaptive Neighbourhoods’ approach^26^, our method progressively refines sample neighbourhoods during training:

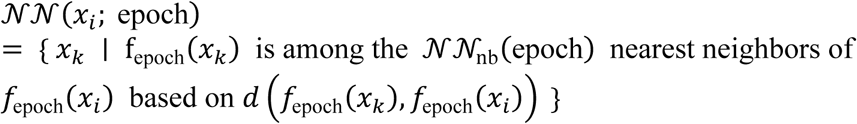

where:

- NNnb is the number of nearest neighbours, which is progressively reduced during training,
- d(*f*(xi), *f*(xk)) is the distance between the embeddings *f*(xi) and *f*(xk).

This allows early learning to capture broad aging patterns, while later training refines local age-related differences, expressed in the adaptive neighbourhood contrastive regression loss:

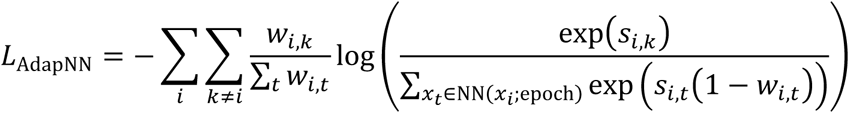

### Evaluation Protocol

Our evaluation strategy comprises three distinct modelling approaches: a kernel-based method, an end-to-end supervised CNN, and a self-supervised method. The first two approaches, kernel-based methods and supervised deep learning, are widely used in brain age prediction studies. Specifically, we apply principal component analysis (PCA) for feature extraction, followed by Gaussian process regression (GP), as a representative kernel method. Additionally, we implement a supervised deep learning model using a ResNet^58^ convolutional neural network (CNN) to learn end-to-end age-relevant features directly from the input data. Beyond these established approaches, we employ the proposed self-supervised learning framework, which uses adaptive neighbourhood sampling to learn meaningful representations for regression tasks. All models were trained using an 80:20 train-test split. Model performance was assessed using the mean absolute error (MAE) on the test set, averaging the results across ten random seeds.

For the kernel-based approach, we employ PCA for feature selection, followed by GPs with linear kernels. The models were trained for 50 epochs using the Adam optimizer, with Gaussian noise serving as an augmentation technique for regularization. Hyperparameter optimization was conducted via random search across 50 iterations for each modality: stiffness-only, damping ratio-only, MRE (damping ratio + stiffness, concatenated and input into PCA), and MRI. The hyperparameters tuned included: Learning rate (lr): [0.0001, 0.001, 0.01, 0.1], PCA components (pca k): [10, 50, 100], Noise standard deviation (std): [0.02, 0.05, 0.1, 0.15, 0.2]. The optimal hyperparameter configurations selected were:

- Stiffness-only: *lr = 0.001, PCA k = 100, noise std = 0.02*
- Damping ratio-only: *lr = 0.001, PCA k = 100, noise std = 0.1*
- MRE: *lr = 0.1, PCA k = 100, noise std = 0.15*
- MRI: *lr = 0.1, PCA k = 100, noise std = 0.2*

For the end-to-end supervised CNN, we selected ResNet-18, the smallest ResNet variant, to align with the dataset scale. The architecture consists of channel dimensions: [input channel, 16, 32, 64, 128]; input channels: 1 for MRI, 2 for MRE; latent head with an embedding dimension of 256; a regression head consisting of a two-layer MLP (256→64→1); ReLU activation throughout the network. Models were trained for 50 epochs using the L1 loss function, a batch size of 32, a learning rate of 5 × 10^<=^and the Adam optimizer. Regularization was implemented using Gaussian noise augmentation, weight decay, and dropout. Hyperparameter tuning was performed via random search across 20 iterations, optimizing: Dropout rate (dropout): [0.1, 0.2], Weight decay: [0.00005, 0.0001], Noise standard deviation (std): [0.02, 0.05, 0.1, 0.15, 0.2]. The optimal hyperparameter configurations selected were:

- Stiffness-only: *dropout = 0.1, weight decay = 0.0001, noise std = 0.2*
- Damping ratio-only: *dropout = 0.1, weight decay = 0.0001, noise std = 0.1*
- MRE: *dropout = 0.1, weight decay = 0.00005, noise std = 0.15*
- MRI: *dropout = 0.1, weight decay = 0.00005, noise std = 0.15*

For the self-supervised approach, the same ResNet-18 architecture is trained in a self-supervised manner using the Adaptive Neighbourhood Contrastive Regression Loss. Following the training of the representations, we employed a Ridge Regression estimator on top of the frozen encoder to predict age. Models were trained for 50 epochs using the Adam optimizer, a batch size of 32, an initial learning rate of 1 × 10^<>^, and a stepwise learning rate decay (reduced by 0.9 every 10 epochs) to calculate the degrees of positiveness of pairs. Hyperparameter tuning, performed via random search across 10 iterations, included: Distance metric: [Manhattan, Euclidean, Similarity], Weight decay: [0.00005, 0.0001], Noise standard deviation (std): [0.1, 0.15, 0.2], Nearest neighbour step size: [1, 2, 5], End nearest neighbour count: [8, 9, 10, 11, 12, 13, 14]. The optimal hyperparameter configurations selected were:

- Stiffness-only: *distance = Euclidean, weight decay = 0.00005, noise std = 0.15, NN step size = 2, end NN count = 8*
- Damping ratio-only: *distance = Euclidean, weight decay = 0.0001, noise std = 0.2, NN step size = 1, end NN count = 14*
- MRE: *distance = Manhattan, weight decay = 0.00005, noise std = 0.1, NN step size = 5, end NN count = 9*
- MRI: *distance = Euclidean, weight decay = 0.00005, noise std = 0.15, NN step size = 1, end NN count = 12*

To facilitate the analysis of the impact of spatial normalisation (Supplementary Fig. S1), we repeated the same evaluation protocol and hyperparameter tuning using the alternative normalisation strategy where images are normalised to mean zero and standard deviation of one on image-level rather than dataset-level. This adjustment allows us to investigate spatial effects of aging independent of global large-scale softening and viscoelasticity trends. The same hyperparameter tuning settings were used as in the primary evaluation. The optimal hyperparameter configurations for GPs were: stiffness-only (lr = 0.01, pca k = 100, noise std = 0.2) and damping ratio-only (lr = 0.001, pca k = 100, noise std = 0.2. For the ResNet-based supervised learning approach, the best configurations were: stiffness-only (dropout = 0.1, weight decay = 0.0001, noise std = 0.02) and damping ratio-only (dropout = 0.1, weight decay = 0.00005, noise std = 0.2). In the self-supervised setting, the optimal model selections were: stiffness-only (distance = Euclidean, weight decay = 0.00005, noise std = 0.1, NN step size = 2, end NN count = 8) and damping ratio-only (distance = Euclidean, weight decay = 0.00005, noise std = 0.15, NN step size = 1, end NN count = 13).

### Saliency Maps

To investigate the spatial importance of brain regions for age prediction, we conducted an occlusion sensitivity analysis^23^ using the trained self-supervised model, the test dataset and the dataset-wide normalisation. This approach allows us to track how mechanical properties influence brain age predictions across different life stages. Test samples were divided into five sub-age groups based on chronological age quantiles. For each group, we systematically occluded 7×7×7 voxel regions by replacing them with zero values. To reduce computational overhead, the superior-most 9 slices along the superior-inferior axis —corresponding to non-brain areas — were excluded from occlusion. After occlusion, the model was used to generate age predictions, and the effect of occlusion was quantified as the difference in mean absolute error (MAE) between occluded and original predictions (delta MAE). Iterating this process for all occlusion regions resulted in a delta MAE matrix of size 13×13×13 (n = 2,197). To focus on relevant variations, values were clipped between the 5th and 95th percentiles. The delta MAE matrix was then resized to match the original image dimensions (91×109×91) using cubic interpolation, with zero padding applied to the excluded areas. Finally, the reconstructed saliency map was normalised by min-max scaling values between 0 and 1.

### Brain Age Gaps

To assess brain age gaps (BAGs) in disease cohorts, models were retrained (with modality-specific best hyperparameters identified in the whole-brain analysis) using only healthy samples while excluding the healthy control subset from the applicable disease cohort. These models were then used to predict ages for both the disease and control cohorts. To correct for systematic biases, a Theil-Sen Regressor was fitted on the control cohort and subsequently applied to the disease cohort. The Brain Age Gap was then computed as the difference between the corrected predicted age and the chronological age for each individual.

### Regional Analysis

To evaluate the regional contributions of brain structures to brain age prediction, brain region-specific masks were applied, and separate models were trained for each region using the best hyperparameters for the specific modality identified in the whole-brain analysis. These models were trained exclusively on healthy samples to establish baseline aging trajectories for different brain regions. For the regional BAG analysis, the same workflow as the whole-brain BAG analysis was followed, where the healthy control cohort was excluded from the training set. The trained region-specific models were subsequently applied to both disease and control cohorts, and the Theil-Sen correction was fitted on the control cohort samples. Individual brain age profiles were computed for each participant, and to facilitate cohort-level interpretations, individual profiles were averaged within each disease group to generate representative cohort-specific brain age profiles.

### Statistics

Statistical analyses were conducted using SciPy^59^ for all comparisons. Data were tested for normality using Shapiro-Wilk test. Depending on the outcome, either a parametric test (paired or independent t-test) or a non-parametric alternative (Wilcoxon signed-rank test or Mann-Whitney U test) was used. For comparisons in Fig. 2, Fig. 5, and Supplementary Fig. S1, paired t-tests or Wilcoxon signed-rank tests were applied. For group comparisons in Fig. 4, independent t-tests or Mann-Whitney U tests were used.

## Acknowledgements

JT acknowledges support from the Gates Cambridge Trust via the Gates Cambridge Scholarship. This project was supported with funding from the Cambridge Centre for Data-Driven Discovery and Accelerate Programme for Scientific Discovery, made possible by a donation from Schmidt Sciences. LVH is supported by the Wellcome Trust (grant number: 226420/Z/22/Z). CJ acknowledges partial support from the National Institutes of Health grants R01-AG058853 and U01-NS112120. CBS acknowledges support from the Philip Leverhulme Prize, the Royal Society Wolfson Fellowship, the EPSRC advanced career fellowship EP/V029428/1, EPSRC grants EP/S026045/1 and EP/T003553/1, EP/N014588/1, EP/T017961/1, the Wellcome Innovator Awards 215733/Z/19/Z and 221633/Z/20/Z, CCMI and the Alan Turing Institute. GSKS acknowledges funding from the Wellcome Trust (065807/Z/01/Z) (203249/Z/16/Z), the UK Medical Research Council (MRC) (MR/K02292X/1), ARUK (ARUK-PG013-14), Michael J Fox Foundation (16238; 022159), and Infinitus China Ltd.

## Supplementary Materials & Methods

**Supplementary Figure 1:**
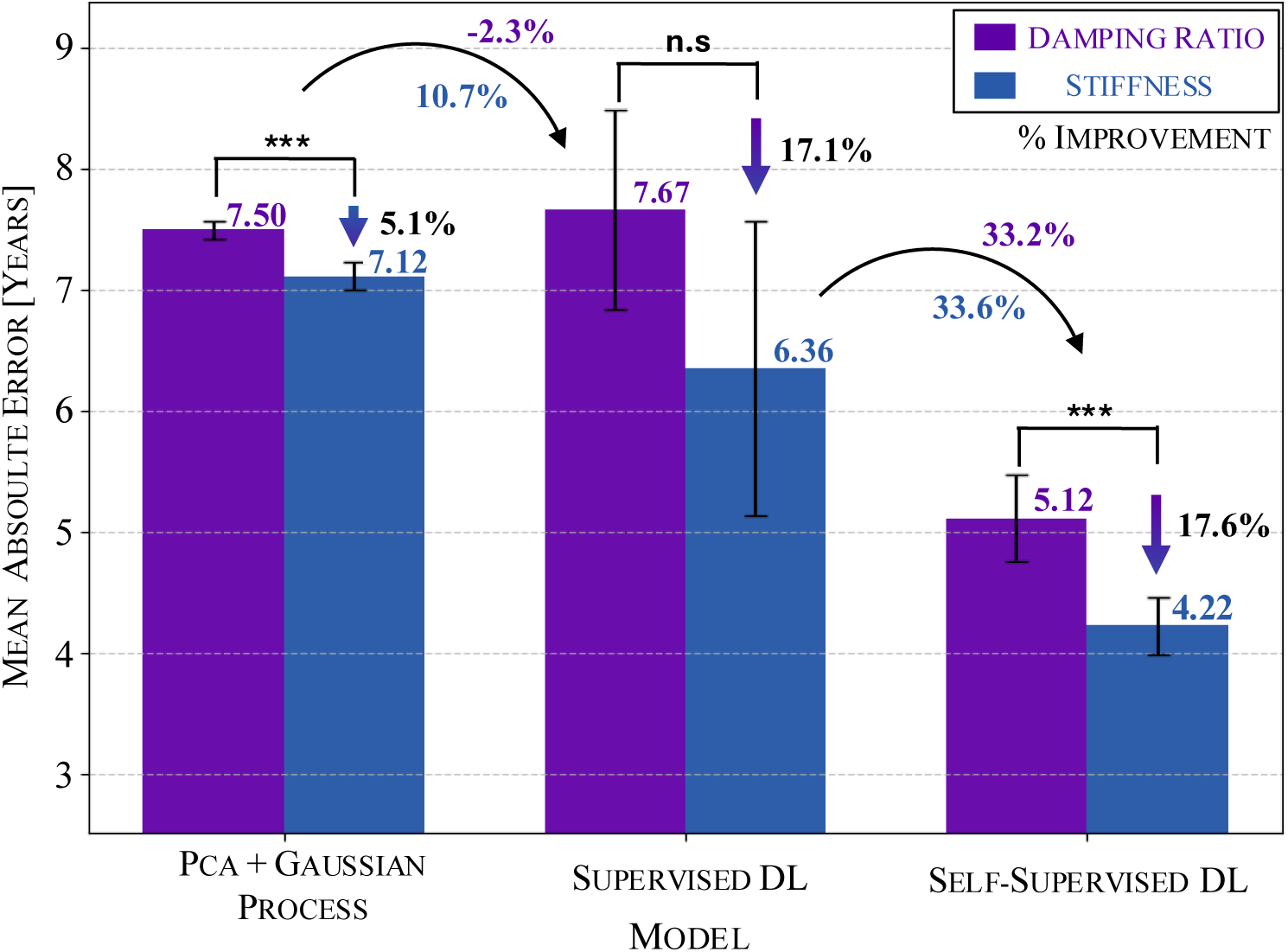
Disentangling Global and Local Aging Effects: Impact of Spatial Normalisation on MRE-Based Predictions. Evaluating spatially normalised mechanical properties highlights the predictive value of localized aging effect. Stiffness proves to be the stronger predictor in all spatially normalised models. To better understand the effects of global versus local aging patterns, we evaluated brain age prediction using spatially normalised MRE scans (see Supplementary Fig. S1). Unlike our primary analysis, which retained global trends in stiffness and damping ratio, this approach normalises each scan to zero mean and unit variance before training. By removing large-scale mechanical aging trends, this normalisation isolates spatially localised aging effects. Under the spatially normalised conditions, stiffness outperforms damping ratio for kernel-based methods. PCA+GPs achieve an MAE of 7.12 years for stiffness, a 5.1% improvement over damping ratio (MAE = 7.50 years). Similarly, in deep learning models, stiffness emerges as the stronger predictor. Supervised learning reduces the MAE to 6.36 years for stiffness compared to 7.67 years for damping ratio, while self-supervised learning further improves stiffness-based predictions to an MAE of 4.22 years. Compared to the previous normalisation method, which preserved global aging trends, supervised and self-supervised deep learning models show slightly higher MAEs, highlighting the predictive value of large-scale mechanical changes. However, this analysis shows that spatially localized distribution changes contain key information for brain age prediction.

**Supplementary Figure 2:**
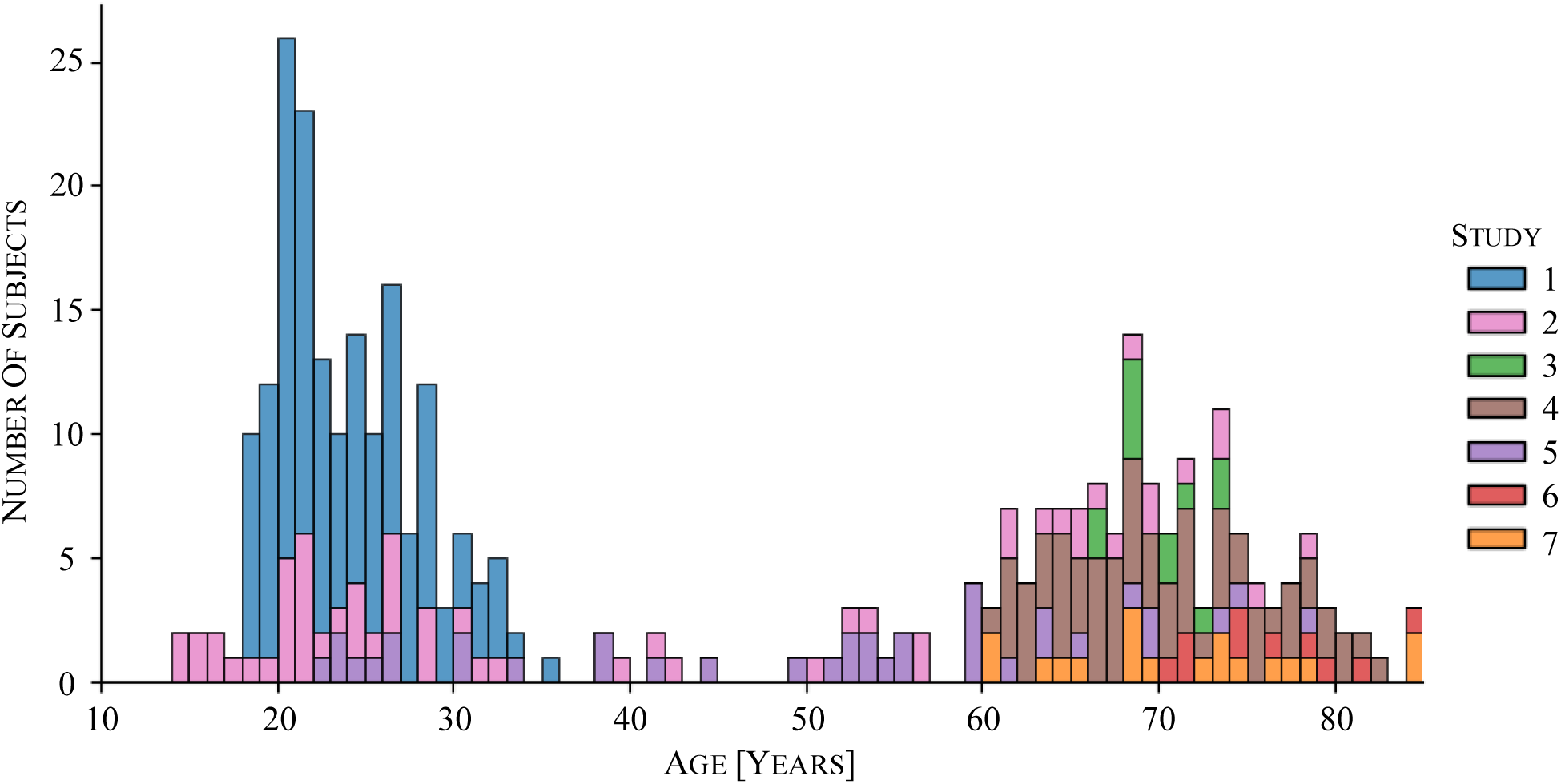
Age Distribution of Pooled Studies of MRE Dataset. Contribution of each study is highlighted in colour. The distribution shows bi-modal characteristics with two predominant age groups among samples. Understanding the age distribution of our dataset is essential for evaluating the generalizability of brain age prediction models. Supplementary Fig. S2 presents the combined age distribution from all pooled studies contributing to the MRE dataset, with each study’s contribution highlighted in different colours. The distribution exhibits a bi-modal pattern, characterised by two predominant age clusters corresponding to younger and older cohorts. This pattern arises due to the inherent limitation of a lack of middle-aged volunteers in clinical trials. To mitigate this imbalance, we employ the adaptive neighbourhood approach, which compensates for dataset non-uniformity through the contrastive regression loss function. This method ensures that age predictions remain robust despite the dataset’s skewed age distribution, enhancing the model’s ability to generalize across age groups.

